# GENI: a web server to identify gene set enrichments in tumor samples

**DOI:** 10.1101/2023.04.05.535584

**Authors:** Arata Hayashi, Shmuel Ruppo, Elisheva E. Heilbrun, Chiara Mazzoni, Sheera Adar, Moran Yassour, Areej Abu Rmaileh, Yoav D. Shaul

## Abstract

The Cancer Genome Atlas (TCGA) and other projects provide informative tumor-associated genomic data for the broad research community. Hence, several useful web-based tools have been generated to ease non-expert users with the analysis and characterization of a specific gene behavior in selected tumors. However, none of the existing tools offer the user the means to evaluate the expression profile of a given gene in the context of the whole transcriptome. Currently, such analyses require prior bioinformatic knowledge and expertise. Therefore, we developed GENI (Gene ENrichment Identifier) as a fast, user-friendly tool to analyze the TCGA expression data for gene set enrichments. GENI analyzes large-scale tumor-associated gene expression datasets and evaluates biological relevance, thus offering researchers a simplified means to analyze cancer patient-derived data.

**Availability and implementation:** GENI is a freely available fast web-based tool developed using the R programming language. It is accessible via the URL: https://yoavshaul-lab.shinyapps.io/gsea-geni/. The complete script can be found at https://github.com/ArataHayashi/GENI-Gene-ENrichment-Identifier.

## 1. Introduction

The TCGA project has revolutionized cancer research by providing a collaborative platform for identifying genomic changes across 33 different cancer types (Liu *et al*., 2018). This open-access database contains vast genomic, transcriptomic, and clinical data generated through various technologies. (Network *et al*., 2013). However, researchers who are non-experts in bioinformatic analyses may find it challenging to navigate and analyze the enormous TCGA database. To address this issue, web-based platforms such as the Genomic Data Commons (GDC) (https://portal.gdc.cancer.gov) (L. *et al*., 2016) and cBioPortal (http://www.cbioportal.org) (Gao *et al*., 2013) have emerged as powerful tools for facilitating the exploration and analysis of the TCGA data, enabling researchers to gain new insights into cancer progression and the identification of potential therapeutic targets.

Despite the availability of various web servers to study selected genes in large-scale databases, understanding the biological meaning of their expressional changes relative to the whole transcriptome remains challenging. Therefore, Gene Set Enrichment Analysis (GSEA) (Subramanian *et al*., 2005) was developed as a computational method to study the behavior of whole gene sets. GSEA provides a comprehensive means to identify pathways or groups that are differentially expressed between various biological conditions.

In this manuscript, we introduce GENI, a web-based platform containing a user-friendly interface for exploring and analyzing TCGA data through GSEA. We aim to provide a comprehensive means for researchers to understand the biological significance of gene expression changes in the context of the whole transcriptome, facilitating the identification of potential therapeutic targets and aiding in cancer research. A more detailed introduction is described in the supplementary materials.

## 2. Methods and software implementation

### Implementation

GENI is a freely available web-based platform developed using the R programming language. It leverages various packages from Bioconductor and CRAN to enable its features and utilizes additional R packages, such as shiny, for data analysis and visualization. Gene set enrichment analysis is performed using the clusterProfiler R package (Yu, 2018).

The expression data used in GENI was obtained from multiple sources, including the TCGA website (https://dcc.icgc.org/releases/PCAWG/), NCBI (https://ftp.ncbi.nlm.nih.gov/gene/DATA/), and from the cBioPortal for Cancer Genomics (http://cbioportal.org) which was accessed using the cBioPortalData R package (Gao 2013; Cerami, 2012). The data were combined and stored on the Shinyapps.io cloud (https://www.shinyapps.io). GENI is designed to be user-friendly, with no login requirements necessary to access its features. A graphical abstract that explains the GENI implementation is illustrated in Figure 1.

**Figure 1:**
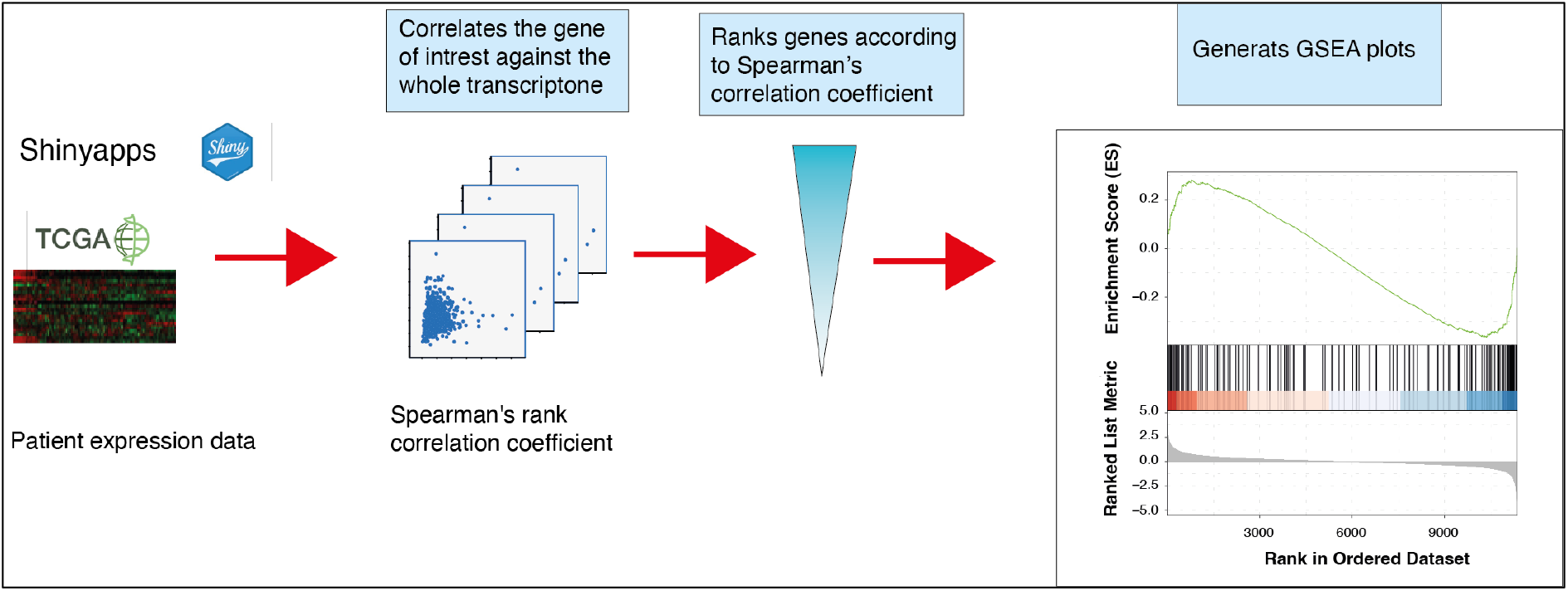
An illustration of the GENI flow. Gene expression data is imported from the Shinyapps.io cloud and used to calculate Spearman’s correlation coefficient for the gene of interest against all protein-coding genes in the transcriptome. The resulting correlation values are then ranked against known gene sets. Finally, GENI produces a table of significantly correlated genes and multiple pathway enrichment graphs of publication quality.

### Workflow overview

The GENI platform provides a user-friendly search function that identifies correlations between a specific gene and the entire transcriptome using GSEA. The search process begins by entering an NCBI gene ID or gene symbol in the “Find your gene” field (Supplementary Fig S1a). The user then selects the desired tissue and study in the “Search tissue and Select your study” field, followed by choosing a gene set from the droplist of gene sets obtained from the Molecular Signatures Database (MSigDB, (Liberzon *et al*., 2011)) (Supplementary Fig S1a). Finally, by clicking “Apply GENI”, Spearman’s correlation coefficients are calculated (rounded to 6 digits) and organized based on correlation values. Moreover, the “Advanced settings” button expands the search options, including permutations, the maximum and minimum number of genes in the gene set, the P-value adjustment, the exponent, and the p-value cut-off (Supplementary Fig S1b).

The results are displayed in a new window as a table and summary dot plot (Supplementary Fig S2a and b). The user can further analyze specific gene sets of interest by clicking the indicated row in the table. This results in the appearance of a GSEA plot (Supplementary Fig S2c), enriched plot values (Supplementary Fig S2d), and a summary table of all the correlations (Supplementary Fig S2e). All of the aforementioned results are presented in a high resolution and can be downloaded in PDF and Excel format through the “Download” button. An example of GENI’s application using the mesenchymal marker CDH2 (N-cadherin) is provided in the supplementary information.

## 3. Comparison with related tools

Web-based platforms such as the Genomic Data Commons (GDC) (https://portal.gdc.cancer.gov) (L. *et al*., 2016) and cBioPortal (http://www.cbioportal.org) (Gao *et al*., 2013) have revolutionized cancer research by providing user-friendly interfaces for researchers to access and analyze the large-scale databases. Furthermore, specialized bioinformatic tools such as Xena (https://xena.ucsc.edu) (Goldman *et al*., 2020) and Gene Expression Profiling Interactive Analysis (GEPIA) (http://gepia.cancer-pku.cn/index.html) (Yang *et al*., 2017) have been developed to analyze TCGA data effectively. Specifically, these web-based tools allow the researchers to perform Kaplan-Meier survival analysis, compare tumor vs. normal within or across tumors, determine the association between increased gene expression and the promotor epigenetic landscape, and create subgroups. These servers provide a platform to effectively navigate and analyze the vast amount of data in the TCGA database, enabling researchers to gain new insights into cancer survival and progression and identify potential therapeutic targets. Overall, the availability of these web-based tools represents a significant advancement in the field of cancer research, making TCGA data more accessible and valuable for researchers.

However, unlike these web tools, GENI offers a unique feature that allows for the comparison of gene expression relative to predefined gene sets. This approach provides a biological context for the analysis of gene expression data and allows for the identification of potential pathways and processes involved in cancer progression. Moreover, the user-friendly interface of GENI makes it easy for researchers to perform complex analyses without requiring extensive bioinformatics expertise. GENI’s unique feature adds another layer of analysis to this wealth of information and provides a valuable resource for the scientific community.

## 4. Conclusion

GENI provides a user-friendly and powerful platform for exploring the TCGA database by allowing researchers to investigate gene expression levels relative to known gene sets. The distinctive feature of this tool is its ability to conduct gene set enrichment analysis, which in turn, enhances the biological relevance of gene expression data analysis. By doing so, it aids in the detection of possible therapeutic targets and brings new perspectives to the study of cancer progression, ultimately benefiting the scientific community. This tool provides a new approach to analyze gene expression data, making it a valuable addition to the field of cancer research. Overall, GENI’s importance lies in its simplicity, biological relevance, and accessibility, making it an attractive tool for researchers studying cancer.

## ACKNOWLEDGEMENT

The authors thank the Shaul Lab team for the helpful discussions about user questions and Emilia Malachi for illustrating the GENI logo.

