## Supplementary data for "GENI: a web server to identify gene set enrichments in tumor samples"

**Example**

To demonstrate GENI's ability to identify biological meaning for specific genes, we used the example of markers for the epithelial-mesenchymal transition (EMT) program. This conserved cellular mechanism plays a significant role in cancer progression, contributing to stem cell properties, drug resistance, immune evasion, and metastasis (Nieto *et al.*, 2016). The induction of the EMT program is orchestrated by signaling pathways in response to extracellular cues such as the transforming growth factor-β (TGFβ) (Katsuno *et al.*, 2019). These changes include loss of cell polarity and cell-to-cell adhesion, along with alterations in the expression levels of cell surface receptors and cytoskeletal reorganization (Brabletz *et al.*, 2021). Additionally, this program induces significant changes in the cell's transcriptomic profile, as the genes associated with the mesenchymal phenotype, such as N-cadherin (CDH2), are upregulated (Bakir *et al.*, 2020). Hence, identifying genes correlating with the EMT hallmark emerged as a promising strategy to identify unknown factors that potentially function in cancer cell aggressiveness.

In this example, we used GENI to search for genes co-expressed with N-cadherin in the breast invasive carcinoma (TCGA, Firehose legacy) dataset. Upon clicking the "Apply GENI" button, the intermediate table (Supplementary Fig S2a) and summary plots (Supplementary Fig S2b) of the GSEA result were displayed in the main panel. From the intermediate table, we selected the "hallmark of epithelial-mesenchymal transition" gene set as it demonstrated the highest normalized enrichment score (NES) value (Supplementary Fig S2b). Upon selecting this gene set, a GSEA plot (Supplementary Fig S2c), a detailed result table (Supplementary Fig S2d), and a gene list for the selected gene set were exhibited (Supplementary Fig S2e). For comparison, we selected the "hallmark of oxidative phosphorylation", as it was the gene set demonstrating the lowest NES value (Supplementary Fig S2f). Our analysis identified that N-cadherin co-expressed genes were strongly correlated with EMT demonstrating the usefulness of GENI in identifying potential factors that function in cancer cell aggressiveness.

**Figure legend**

**Figure S1: search panel.** (a) A screenshot of the search panel where CDH2 in breast cancer samples is demonstrated as an example. (b) A screenshot of the advanced search option.

**Figure S2: CDH2 expression correlates with EMT markers in breast cancer patients.** (a) a screenshot of the summary table. (b) Summary dot plot of upregulated and downregulated

gene sets of CDH2 correlated genes colored by FDR, sized by gene set size, then sorted in ascending order of p-value. (c) GSEA plot of the Hallmark of EMT shows a positive and significant correlation of CDH2 with the EMT markers in breast cancer patients. NES: Normalized enrichment score FWER: Family-wise error rate FDR: False discovery rate
(d) Screenshot of the detailed result of the Hallmark of the EMT GSEA plot. (e) A screenshot of the EMT gene list and Spearman's rank correlation coefficient with CDH2 expression. (f) GSEA plot of Hallmark of Oxidative Phosphorylation (OXPHOS) shows a negative correlation with CDH2 expression in breast cancer patients. NES: Normalized enrichment score FWER: Family-wise error rate FDR: False discovery rate
