## Supplementary figures for "GENI: a web server to identify gene set enrichments in tumor samples"

(a)

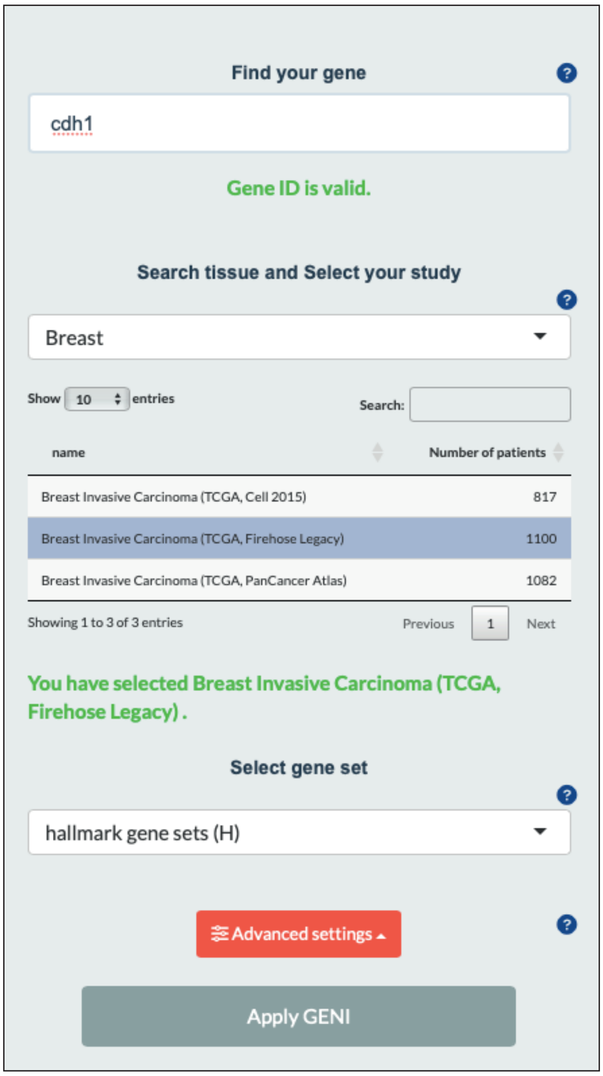

Find your gene
cdh1
Gene ID is valid.
Search tissue and Select your study
Breast
Show 10 entries
Search:
name
Number of patients
Breast Invasive Carcinoma (TCGA, Cell 2015) 817
Breast Invasive Carcinoma (TCGA, Firehose Legacy) 1100
Breast Invasive Carcinoma (TCGA, PanCancer Atlas) 1082
Showing 1 to 3 of 3 entries
Previous 1 Next
You have selected Breast Invasive Carcinoma (TCGA, Firehose Legacy) .
Select gene set
hallmark gene sets (H)
Advanced settings
Apply GENI


(b)

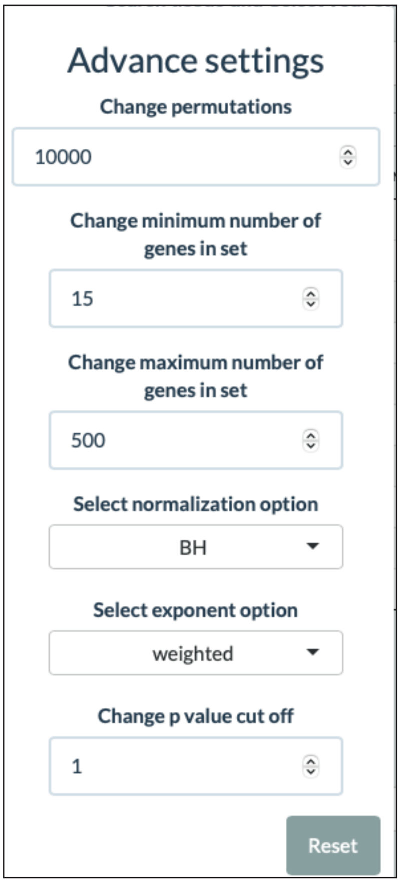

Advance settings
Change permutations
10000
Change minimum number of genes in set
15
Change maximum number of genes in set
500
Select normalization option
BH
Select exponent option
weighted
Change p value cut off
1
Reset


### Figure S1

(a)

#### Choose the Gene Set to get details

Show 10 entries Search:

| Download | GeneSet | Size | NES | NOM p-val | FWER p-val | FDR q-val |
|---|---|---|---|---|---|---|
| ☐ | HALLMARK_EPITHELIAL_MESENCHYMAL_TRANSITION | 197 | 3.33639718308712 | 3.3704451954154e-63 | 1.6852225977077e-61 | 4.61218816214739e-62 |
| ☐ | HALLMARK_ANGIOGENESIS | 36 | 2.42737414732837 | 1.26814131922894e-9 | 6.3407065961447e-9 | 1.73535127894487e-9 |
| ☐ | HALLMARK_TGF_BETA_SIGNALING | 54 | 2.40300733105737 | 3.67707466661517e-10 | 2.04281925923065e-9 | 5.59087376210494e-10 |
| ☐ | HALLMARK_UV_RESPONSE_DN | 144 | 2.3442970948624 | 1.04640908123011e-13 | 1.30801135153763e-12 | 3.57982054105037e-13 |
| ☐ | HALLMARK_KRAS_SIGNALING_UP | 200 | 2.25945185867619 | 3.44101145607886e-14 | 5.73501909346477e-13 | 1.56958417294825e-13 |
| ☐ | HALLMARK_G2M_CHECKPOINT | 200 | 2.23004013485713 | 1.55477357655824e-13 | 1.55477357655824e-12 | 4.25516978847519e-13 |
| ☐ | HALLMARK_APICAL_JUNCTION | 200 | 2.18800585018952 | 1.29577535803468e-12 | 1.07981279836223e-11 | 2.95527713235979e-12 |
| ☐ | HALLMARK_INFLAMMATORY_RESPONSE | 200 | 2.15675232344934 | 6.78783857557657e-12 | 4.84845612541184e-11 | 1.32694588695482e-11 |
| ☐ | HALLMARK_MITOTIC_SPINDLE | 199 | 2.13448187598583 | 1.38049993123593e-11 | 8.62812070224564e-11 | 2.36138040271986e-11 |
| ☐ | HALLMARK_COAGULATION | 138 | 2.06329710104318 | 1.02519555338073e-8 | 3.94305982069511e-8 | 1.07915321408498e-8 |

Showing 1 to 10 of 50 entries Previous 1 2 3 4 5 Next

(b)

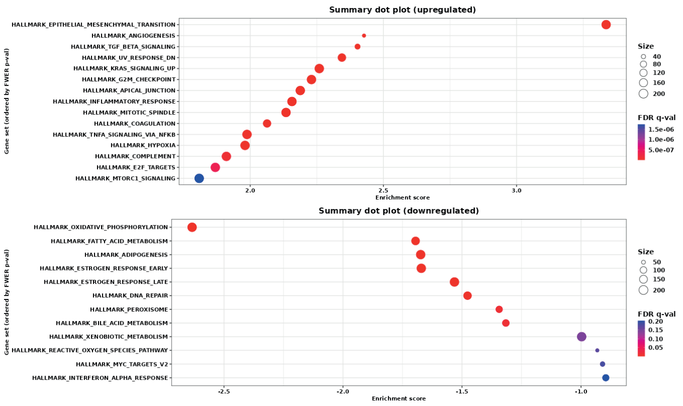


(c)

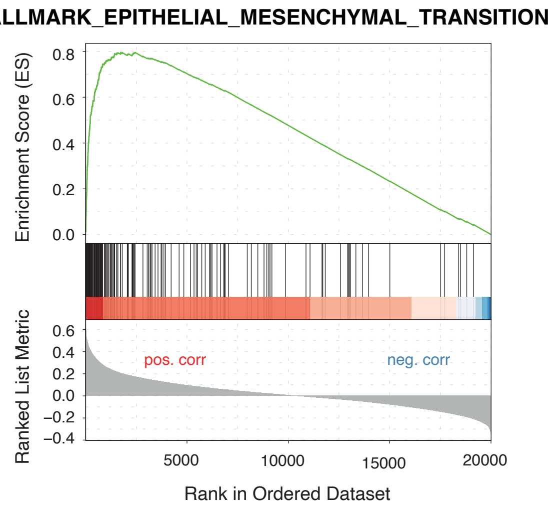


(d)

| Name | Value |
|---|---|
| Enrichment Score (ES) | 0.7954 |
| Normalized Enrichment Score (NES) | 3.3364 |
| Nominal p-value | 3.370e-63 |
| FWER p-value | 1.685e-61 |
| FDR q-value | 4.612e-62 |

(e)

Show 10 entries Search:

| Symbol | Entrez gene ID | Rank Metric Score | Core Enrichment |
|---|---|---|---|
| FN1 | 2335 | 0.640239654533688 | Yes |
| COL11A1 | 1301 | 0.587180934380594 | Yes |
| INHBA | 3624 | 0.583802665615582 | Yes |
| THBS2 | 7058 | 0.578346150199702 | Yes |
| CALD1 | 800 | 0.561598211452162 | Yes |
| LOXL2 | 4017 | 0.558487256206945 | Yes |
| LOX | 4015 | 0.555544482668819 | Yes |
| GREM1 | 26585 | 0.55504449753124 | Yes |
| CDH11 | 1009 | 0.549207823759469 | Yes |
| COL5A2 | 1290 | 0.547781985126619 | Yes |

Showing 1 to 10 of 197 entries

Previous 1 2 3 4 5 ... 20

(f)

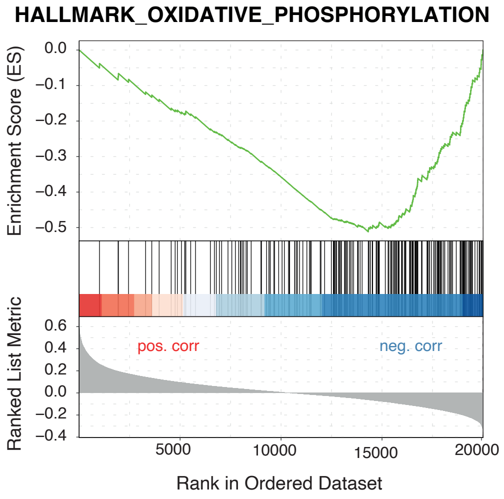


### Figure S2
